# Identifying and prioritizing demersal fisheries restricted areas based on combined ecological and fisheries criteria: the western Mediterranean

**DOI:** 10.1101/2023.02.02.526784

**Authors:** Miquel Ortega, María D. Castro-Cadenas, Jeroen Steenbeek, Marta Coll

## Abstract

The western Mediterranean basin is a high marine biodiversity area under severe pressure by changing climate and intense human activities. Beyond national jurisdictions, international institutions such as the General Fisheries Commission for the Mediterranean (GFCM) work towards canalizing a regional consensus that fishing practices should evolve to better support the 2030 Agenda for Sustainable Development. In this context, Fisheries Restricted Areas (FRA) are proposed as effective management measures to contribute towards increasing fisheries sustainability in the region that can be considered, under some conditions, as Nature-based Solutions (NbS); however, how to operationalize their framework remains unclear. In this study, based on combined ecological and fisheries criteria, we identify and prioritize six potential priority areas for management (PAMs) in the western Mediterranean Sea. They are specifically aimed at the protection and recovery of Essential Fish Habitats and the conservation of Vulnerable Marine Ecosystems, whilst requiring limited adaptation of fisheries practices due to their relative low fishing pressure. We compare the identified areas to those that are currently under protection, and to areas that have been proposed for protection at the GFCM. Our results show that the FRAs and other spatial management measures introduced in the last years marginally contribute to the protection PAMs in the western Mediterranean region. However, the adoption of FRAs that are currently under discussion at the GFCM could contribute significantly to improve the situation. FRAs could also contribute to operationalize NbS in the western Mediterranean Sea when properly designed and implemented.

**Highlights:**

- Based on combined ecological and fisheries criteria, six priority areas for management (PAMs) in the western Mediterranean Sea have being identified, with multiple ecological values and relative low trawling.
- Current spatial management measures implemented have little contribution on PAMs protection.
- Fisheries Restricted Areas currently under discussion at the GFCM can significantly increase the protection level of high priority PAMs.

## 1. Introduction

The Mediterranean basin is a biodiversity hotspot with high diversity and endemism of flora and fauna (Coll et al., 2010; UNEP/MAP-Plan Bleu, 2020). Nevertheless, severe pressures such as fishing, pollution, coastal uses, maritime transport, climate change and the introduction of invasive species are changing its characteristics (Coll et al., 2012, 2015; Micheli et al., 2013a; Piroddi et al., 2020) and hinder the opportunity to achieve a good environmental status of the marine ecosystem and the Sustainable Development Goal (SDG) 14 *“Conserve and sustainably use the oceans, seas and marine resources”* (United Nations, 2015) in the region.

Historically, fisheries have had an important socio-economic role in the western Mediterranean basin. According to the General Fisheries Commission for the Mediterranean (GFCM), the fishing fleet of the western Mediterranean consists of 15.552 vessels, and provides 241.626 tons of catches annually (FAO, 2022d). While its direct contribution to revenue is small in comparison with other economic sectors (with a total to 977 million USD in 2020), fishing remains a relevant socio-economic and cultural activity associated with more than 42.800 jobs (FAO, 2022d).

Fishing has become economically less relevant in the past twenty years, and faces important challenges to ensure long-term sustainable well-being of coastal communities. Exploitation rates are still too high, with 26 of the 29 evaluated stocks in the western Mediterranean Sea requiring reduced fishing mortalities (FAO, 2022b). Moreover, current fishing practices have cascading impacts on marine food webs (e.g., Coll et al., 2009, 2021; Colloca et al., 2017) and seafloor habitats (Bastari et al., 2022) that hamper ecological recovery.

There is a regional consensus that fishing practices should evolve to better support the 2030 Agenda for Sustainable Development (United Nations, 2015), a goal that is reflected in the GFCM 2030 Strategy (FAO, 2021a). Potential tools to manage fisheries include Area-Based Fisheries Management measures (FAO, 2021a; Petza et al., 2021), intended to protect key elements of marine ecosystems to contribute to the recovery of habitats and species. Especially, the use of marine spatial areas to protect Vulnerable Marine Ecosystems (VME) and Essential Fish Habitats (EFH), when integrated in an ecosystem-based approach, have proven to be effective for managing fisheries and to improve ecosystem health (McConnaughey et al., 2020; Smith et al., 2006). If properly designed and implemented, they can be considered Nature-based Solutions (NbS; Cohen-Shacham et al., 2019) by simultaneously addressing societal problems such as “climate change mitigation and adaptation”, “economical & society development”, “food security”, and “environmental degradation & biodiversity loss” (Riisager-Simonsen et al., 2022). They can also be considered Other Effective Area-Based Conservation Measures (OECMs) as they can contribute to achieve positive long-term outcomes for protecting biodiversity and associated ecosystem services and functions, and other locally relevant social or economic values (CBD, 2018).

Vulnerable Marine Ecosystems (VMEs) are benthic groups of species, communities, or habitats that may be vulnerable to impacts from fishing activities (FAO, 2009). They are considered hotspots of biodiversity and ecosystem functioning and they provide habitat, nursery areas and feeding grounds for marine organisms. They are fundamental for maintaining healthy ecosystems, as they perform a wide range of ecosystem services (e.g., storing carbon, filtering and supporting food provisioning) and some VME animals have potential for bio-discovery. Overall, they have low reproduction and growth rates, and thus can only sustain low exploitation rates while recovery can be slow and uncertain. Essential Fish Habitats (EFH) include all types of aquatic habitat where fish spawn, breed, feed or mature, such as wetlands, coral reefs, seagrasses and rivers. In the marine environment, coral gardens, kelp forests, sponge beds, seagrass meadows and submarine canyons represent EFH essential for fish survival. They are also very sensitive to human activities, mostly to bottom trawling and dredging, and their protection can contribute to the rebuilding and sustainably exploiting fish stocks.

VMEs and EFHs are usually detected and monitored through *in situ* observations or through scientific campaigns (DeLaHoz et al., 2018; Druon et al., 2015; Giannoulaki et al., 2013a; Paradinas et al., 2015; Pennino et al., 2020), non-invasive observation technologies (Chimienti et al., 2021; Giakoumi et al., 2013), and local ecological knowledge (Bastari et al., 2022). In addition, statistical modelling is frequently used to predict the distribution of VMEs and EFHs, recently even considering climate change projections (Bleuel et al., 2021; Izquierdo et al., 2021; Morato et al., 2020).

The use of spatial management tools to protect key biodiversity elements and enhance fishing in the Mediterranean is not new. There are several national, regional and multi-national political commitments, processes and institutions in place towards the implementation of spatial management (Claudet et al., 2020; Micheli et al., 2013b). These include, among others, the designation of marine protected areas, Natura 2000 sites, Specially Protected Areas of Mediterranean Importance (SPAMIs), Important Birds Areas (IBAs), as well as other national specific legal measures that restrict areas to fishing.

In addition, the GFCM has had a major role in the development of spatial management measures through mandatory decisions approved by its members. Currently, ten Fisheries Restricted Areas (FRAs) in non-territorial waters have already been established and implemented by the GFCM (FAO, 2022), with two (the Gulf of Lions FRA and the deep-sea FRA) that cover part of the western Mediterranean (Figure 1). Moreover, the new GFCM 2030 strategy targets the extension of Fisheries Restricted Areas as one of its objectives over the period 2022-2030 (FAO, 2021a).

**Figure 1.**
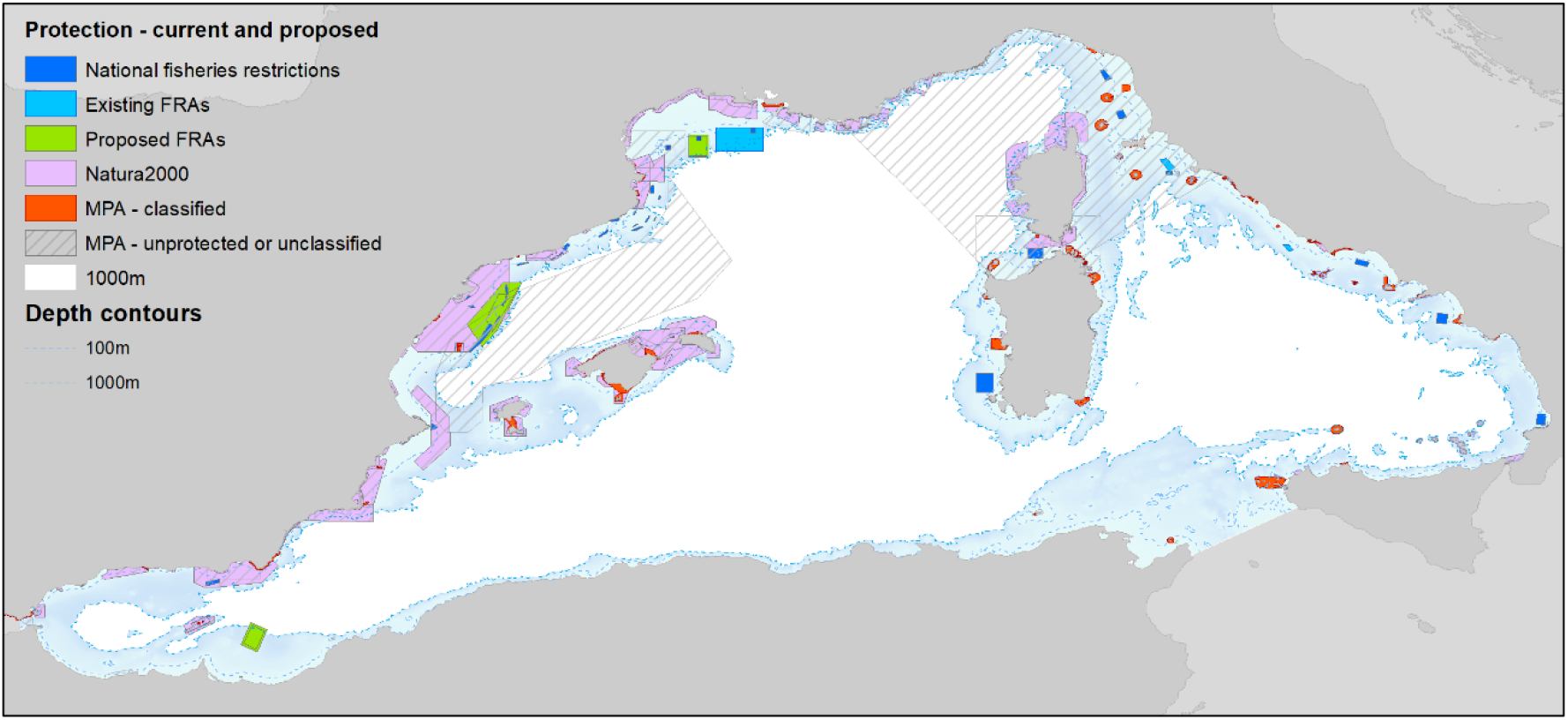
Current and potential protection areas in the western Mediterranean Sea.

The European Union (EU) is another relevant political actor supporting the expansion of spatial measures in the Mediterranean. For instance, provisions for spatial fisheries measures are included in the EU Multiannual plan for demersal fisheries in the western Mediterranean (Regulation (EU) 2019/1022). The EU also promotes them through the proposal of GFCM recommendations, public funding through the European Maritime, Fisheries and Aquaculture Fund (EMFAF), and through political strategies such as the EU Biodiversity Strategy for 2030. The EU has also expressed support for spatial fisheries measures in multiple high-level multilateral political declarations and the European Commission has included them as key elements in the new Nature Restauration Regulation, currently under negotiation with the European Council and the European Parliament.

Nevertheless, scientific, socio-economic and political challenges remain in the definition, prioritization and implementation of these areas. While there is a consensus that these steps should be backed by clear scientific information (Micheli, et al., 2013a), proposed methods are still evolving (Scientific Technical and Economic Committee for Fisheries, 2022a), while the current contribution of spatial fishing measures to sustainability is not systematically assessed it is presumed modest (Claudet et al., 2020; Corrales et al., 2020; Vilas et al., 2021).

Building on the concept of “Low hanging fruits for conservation” (Coll et al., 2015), the aim of this paper is to support the identification and prioritization of permanent Fisheries Restricted Areas in areas where conservation is both feasible and ecologically relevant. We examined non-territorial seas of the western Mediterranean Sea using a combined ecological and fisheries approach based on the best public spatial data available. We first reviewed, mapped and harmonized data available regarding VMEs and EFHs in the western Mediterranean Sea. We then overlaid the different ecological datasets to identify key areas with multiple ecological relevance. We identified which ecological valuable areas were least fished, as fishing adaptation is presumably easier implemented in low fished areas. Finally, we evaluated to which extent these high-value ecological and less fished areas are included in current and discussed fisheries management, considering two distinct political approaches to the selection of spatial management areas: (i) the “current scenario”, which is based on the fishing restricted areas to trawling implemented by the end of 2022, in addition to national designated areas, and (ii) the “discussed scenario”, that covers all potential new fisheries restricted areas currently under discussion in the GFCM. We discussed our results, the relationship between the existing and under discussion protection areas and the identified priority areas, and the potentials and challenges of the approach. We conclude with some suggestions for next steps in terms of sustainable fisheries management and the implementation of Nature-based Solutions in the western Mediterranean Sea.

## 1. Methods

### 2.1. Study area

Our analysis covers the non-territorial waters of the western Mediterranean Sea (FAO subarea 37.1, Figure 1). The western basin (^~^ 846,000 km2, 0-3600 m depth) includes waters from European (Spain, France, Italy, Malta) and African (Morocco, Algeria and Tunis) countries, and spans five marginal seas: the Thyrrenian Sea, the Balearic Sea, the Sea of Sardinia, the Ligurian Sea and the Alboran Sea.

The region is the most productive of the entire Mediterranean Sea; especially the northwestern region and the continental shelves are associated with large rivers and deltas (Rhone and Ebro Delta) and counter-clock wise water circulation (Bosc et al., 2004). It hosts a high diversity of Mediterranean habitats and species, including endemic and at-risk species of seabirds, marine turtles, marine mammals, chondrichthyans, finfish, invertebrates and primary producers (Coll et al., 2012; Coll et al., 2010).

The region is one of the principal maritime corridors in the world and is the gateway to Africa for European countries (Katsanevakis et al., 2015). Maritime activities and a highly urbanized and industrialized coastline threaten the ecosystem through air and water pollution, waste generation, resource degradation and depletion, among others (Coll et al., 2012; Katsanevakis et al., 2015; Micheli et al., 2013a). Climate change poses increasing impacts to habitats and resources, and is expected to intensify more rapidly than the average global mean (Calvo et al., 2011; Garrabou et al., 2019; Marbà et al., 2015; Moatti & Thiébault, 2016; Salat et al., 2019). Additionally, overexploitation of fishing resources is further degrading the ecosystem, while current conservation measures are insufficient to halt biodiversity loss and declining ecosystem health (Claudet et al., 2020).

### 2.2. Spatial data

#### 2.2.1. Ecological data

The ecological analysis was based on the geographical distributions of EFH species and VME habitats for 2012-2020.

Key EFH species included commercial and vulnerable demersal species prioritized by the GFCM (GFCM, 2021a; group 1 and group 2 species), complemented with species included in the European Union Multiannual management plan for demersal stocks in the western Mediterranean Sea. Only species for which geographic distribution data was freely available were included in the analyses (Table 1 and Figures S1 and S2).

**Table 1.** EFH data used for the analysis.

| Species included | Notes | Reference |
| --- | --- | --- |
| **EFHs of key commercial species** |  |  |
| Spawning areas of fish (*Merluccius merluccius*, *Mullus barbatus*, *Pagellus erythrinus*), cephalopods (*Eledone cirrhosa*, *Illex coindetii*), crustaceans (*Aristaeomorpha foliacea*, *Parapenaeus longirostris*, *Nephrops norvegicus*, *Aristeus antennatus*) |  | (Giannoulaki et al., 2013b); MEDISEH project<sup>1</sup> |
| Nursery areas of fish (*Merluccius merluccius*, *Pagellus erythrinus*), cephalopods (*Eledone cirrhosa*, *Illex coindetii*), crustaceans (*Aristaeomorpha foliacea*, *Parapenaeus longirostris*, *Nephrops norvegicus*) |  | (Giannoulaki et al., 2013b); MEDISEH project<sup>1</sup> |
| Nursery areas of hake (*Merluccius merluccius*) | 2015-2018, with >75% suitability | (Druon et al., 2015). Source: https://data.jrc.ec.europa.eu/dataset/3c1044f7-0728-47d1-ac7d-78505082c38d |
| **EFHs of key vulnerable species** |  |  |
| Spawning areas of elasmobranchs (*Raja clavata*, *Galeus melastomus*) |  | (Giannoulaki et al., 2013b); MEDISEH project<sup>1</sup> |
| Nursery areas of elasmobranchs (*Raja clavata*, *Galeus melastomus*) |  | (Giannoulaki et al., 2013b); MEDISEH project<sup>1</sup> |

<sup>1</sup> https://imbriw.hcmr.gr/mediseh/

VME habitat data was obtained from a range of published and open data sources (Table 2). VMEs are generally unevenly distributed. *Isidella elongata* is mostly found in the Gulf of Lions with relevant patches in the Catalan Sea, Corsica island and western Sicily, while maërl beds are distributed in multiple scattered areas of the Tyrrhenian and Ligurian Sea including Sardinia; and coralligenous can be found in most of the areas as *Posidonia oceanica* beds, but concentrated in shallower waters (Figure S3).

**Table 2.**
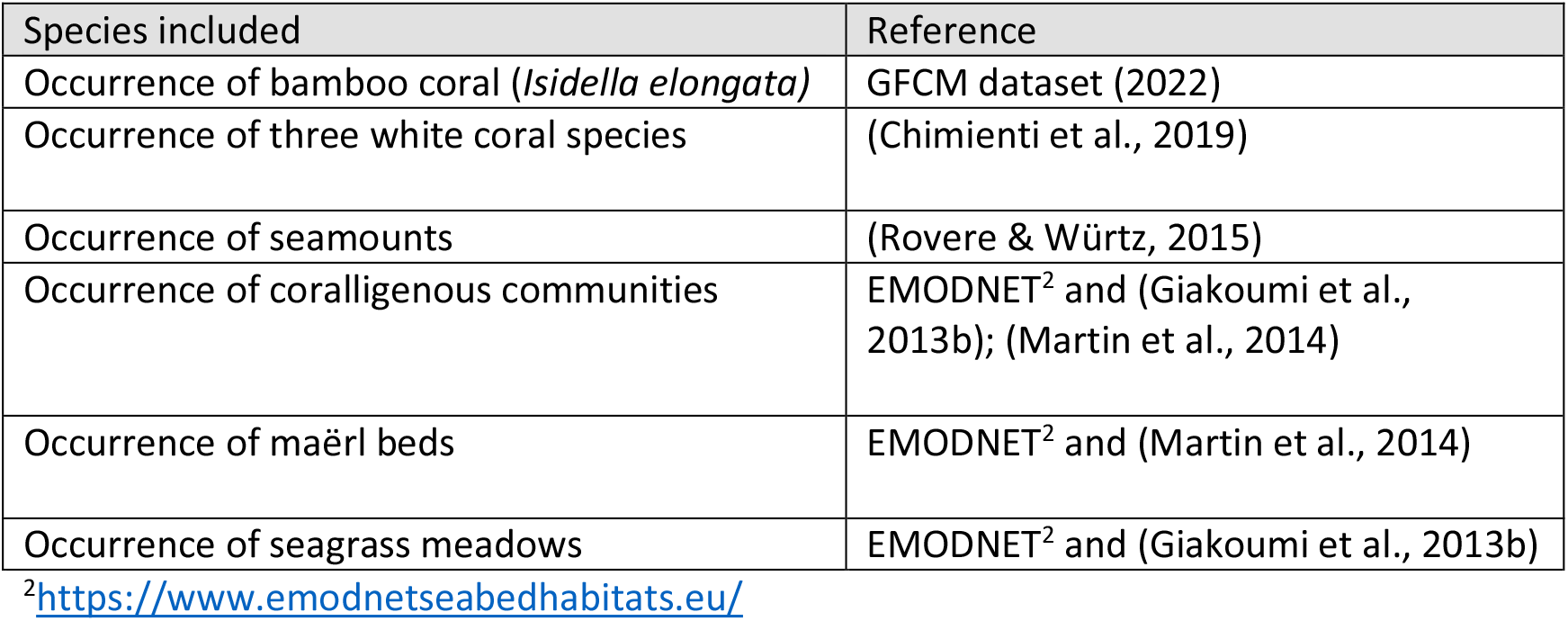
VME data used for the analysis.

#### 2.2.2. Fisheries data

The fisheries analysis was based on trawling fishing effort provided by Global Fish Watch (GFW, 2022) coming from the Automatic Identification System (AIS) ^1^. We selected the year 2019, the most recent and complete annual data available prior to COVID disruptions. Trawling data was chosen due to the severe impacts of trawl nets on the benthic ecosystem, especially on VMEs (UNEP/MAP-Plan Bleu, 2020).

#### 2.2.3. Management data

The management analysis focused on all spatial existing and projected measures that prohibit trawling in some form, including both highly protected and no-take MPAs, FRAs, and national restricted fishing areas, as approved by the end of 2022 or proposed as potential new areas (Figure 1 and Table 3).

**Table 3.** Current and proposed management measures considered in this study for the western Mediterranean Sea.

| Management measures | Notes | Reference |
| --- | --- | --- |
| *Currently in place* |  |  |
| Nationally-designated highly protected MPAs | In territorial waters | (*Protected Planet*, 2022); (Claudet et al., 2020) |
| No-take sub-area of the Gulf of Lion FRA | Closed to trawling since 2020. | (Recommendation GFCM/44/2021/5) |
| Deep sea 1000 m Fisheries Restricted Area | Closed to trawling since 2005. | (Recommendation GFCM/29/2005/1) |
| National designated restricted fishing areas: French permanent closures in the Gulf of Lions area GSA7 and Corsica GSA8; areas in the Spanish | Established in 2020 | (Orden APA/423/2020), (Decreto direttoriale di attuazione del l'art.6, comma 1 del D.M. N°13128), (Arrêté Du 20 Décembre 2019 Portant |

<sup>1</sup> https://globalfishingwatch.org/

|  |  |  |
| --- | --- | --- |
| coast in GSA 1, 2, 5 and 6, and in Italy in GSA 9, 10 and 11 |  | Modification de l'arrêté du 28 Février 2013 portant l'adoption d'un Plan de Gestion pour la pêche professionnelle au chalut en mer Méditerranée par les navires battant pavillon français, 2019) |
| *FRA proposals under discussion by the GFCM* |  |  |
| Expansion of the deep-sea FRA 1000 meters trawl ban to 600-800 meters | Mediterranean region | (FAO, 2022b) |
| Three proposed FRAs: the "Ebro Delta Margin FRA", the "Gulf of Lions Martí and Sete Canyons FRA", and the "Alboran Sea Cabliers coral mound province FRA" | In Spain, France, Morocco and Algeria | (FAO, 2021b), (FAO, 2022a) |

### 2.3. Multi-Ecological Value Areas (MEVAs), Fishing pressure hotspots (FPH) and Priority Areas for Management (PAMs)

Some data processing was needed to make data suitable for spatial analysis. VME point-source and line data was expanded to polygons with a 2500m geographic buffer to provide a reasonable spread of VME presence. VME probability of occurrence polygons were included where probabilities exceeded 75%.

Each EFH and VME data layer, in polygon format, was converted to a spatial grid with a cell size of 0.01 x 0.01 decimal degrees, where each cell recorded the absence or presence of a given EFH or VME feature. Using equal weight, thus not favoring any particular EFH and VME, the number of presences of EFH and VME features were summed per cell to establish what we call “Multi-Ecological Value Area” (MEVA) maps. In order to facilitate the interpretation of results, overlapping feature counts were classified in quartiles (1 feature, 2-3 features, 4-6 features and more than 6 features).

GFW-derived trawl hours for 2019 were classified into quartiles, and were reclassified to an inversed fishing pressure intensity index ranging from 1 to 4, where low trawl intensities scored highest (and high trawl intensities scored lowest), creating what we call “Fishing Pressure Hotspots” (FPH) maps. FPH maps were created for the entire western Mediterranean, and for the Exclusive Economic Zones (EEZs) of France, Italy and Spain scaled to the range of trawl hours in each respective EEZ.

MEVAs and FPH maps were multiplied to obtain what we refer to as the “Priority Areas for Management” (PAM) maps, with values ranging from 0 to 16, where higher values correspond to high MEVA overlap with low fishing pressure. To facilitate the interpretation of results, the PAM maps were reclassified into quartiles according to their potential to become conservation areas for the protection of demersal communities: very high priority, high priority, medium priority and low priority.

PAMs were produced for the wider western Mediterranean to provide a regional perspective, and for the EEZs of France, Italy and Spain to provide national perspectives.

### 2.4. PAM / protection overlap scenarios

Finally, we calculated the percentages that the PAM priority areas are covered by fisheries restricted areas. We considered four different protection scenarios (Table 3):

1. The “current” scenario, which includes the permanent FRAs and national fishing restricted areas including highly protected MPAs, as implemented by the end of 2022. The areas include among others the recently implemented French permanent closures in the Gulf of Lions area GSA7 and Corsica GSA8^2^, the new Spanish restricted areas in GSA 2, 5 and 6^3^, and those introduced by Italy in GSA 9, 10 and 11^4^, and the existing deep sea 1000 meters FRA (Figure 2 and Table 3).
2. The “discussed” scenario, which extends the current scenario with new areas that have been endorsed by the GFCM Scientific Advisory Committee (SAC) or are under evaluation by the GFCM (Figure 2): the establishment of the “Ebro Delta Margin FRA” (FAO, 2021b), the “Gulf of Lions Martí and Sète Canyons FRA” (FAO, 2022a) and the “Alboran Sea Cabliers coral mound province FRA” (FAO, 2022a).
3. The “800m” scenario, which extends the discussed scenario (point 2 above) by expanding the current deep sea 1000 meters FRA to 800 meters.
4. The “600m” scenario, which extends the discussed scenario (point 2 above) by expanding the deep-sea FRA to 600 meters (FAO, 2022b).

**Figure 2.**
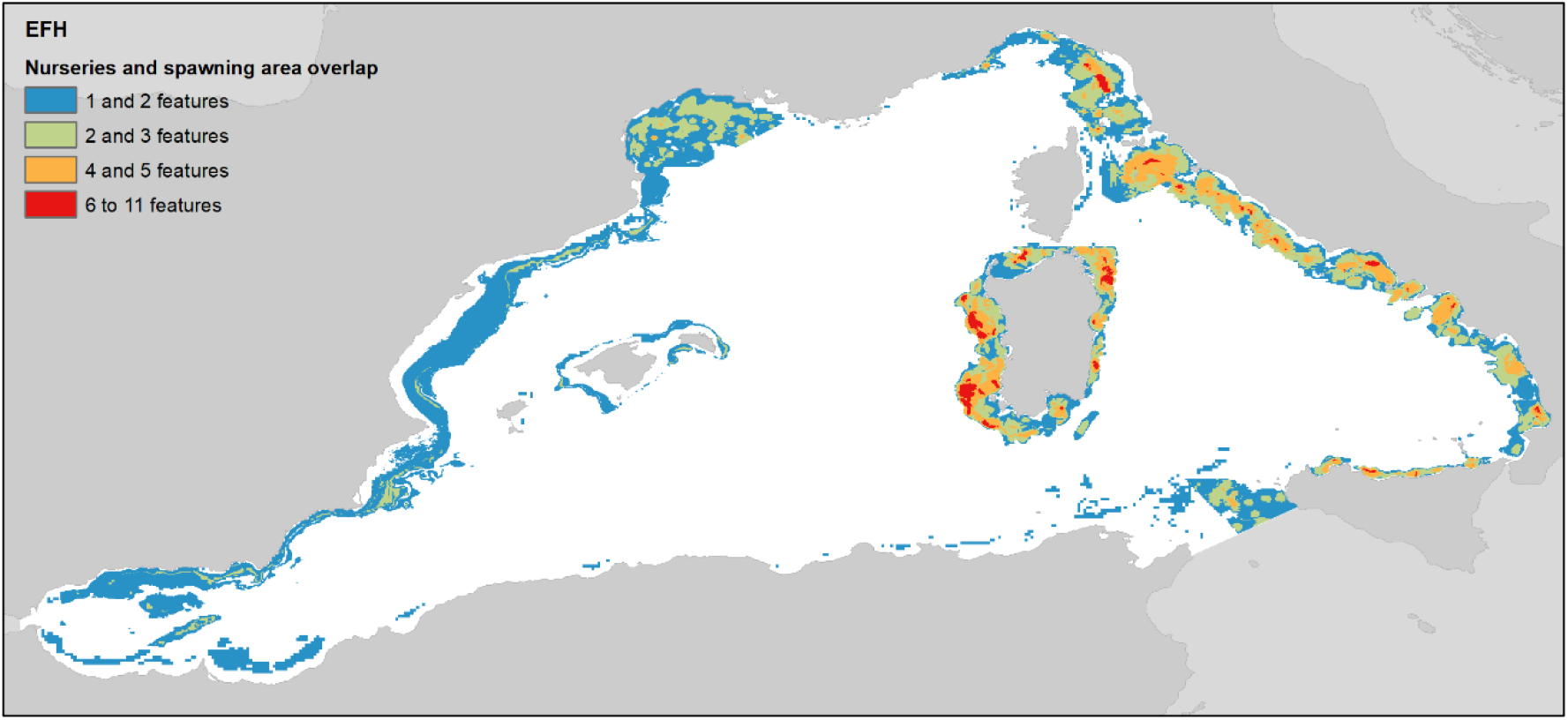
Overlap count of nurseries and spawning areas in the western Mediterranean Sea.

We performed this calculation for the entire western Mediterranean and for the EEZs of France, Italy and Spain. Calculations excluded territorial seas, within 12 nautical miles from the coast, which are considered sovereign territory of states according to United Nations Convention on the Law of the Sea (United Nations, 1982).

## 3. Results

### 3.1. Multi-Ecological Value Areas (MEVAs)

The highest EFH overlap are found around Sardinia, along the eastern Italian coast and in the Gulf of Lions, with a few additional hot spots in Spanish waters (Figure 2). Detailed information of the distribution of EFHs can be found in the Supplementary Material (Figure S4).

As shown in Figure 3 the highest concentrations of EFH and VME overlap are located around Sardinia (Area 1), the eastern part of the Ligurian Sea, the eastern and southern part of the Tyrrhenian Sea (Area 2) and the Gulf of Lions (Area 3). Other areas of interest are located in the western part of Sicily (Area 4), the Ebro Delta (Area 5), the Murcia region (Area 6) and in the Alboran Sea (Area 7). Much lower levels of overlap are found along the Northern African coast and near the French / Italian border.

**Figure 3.**
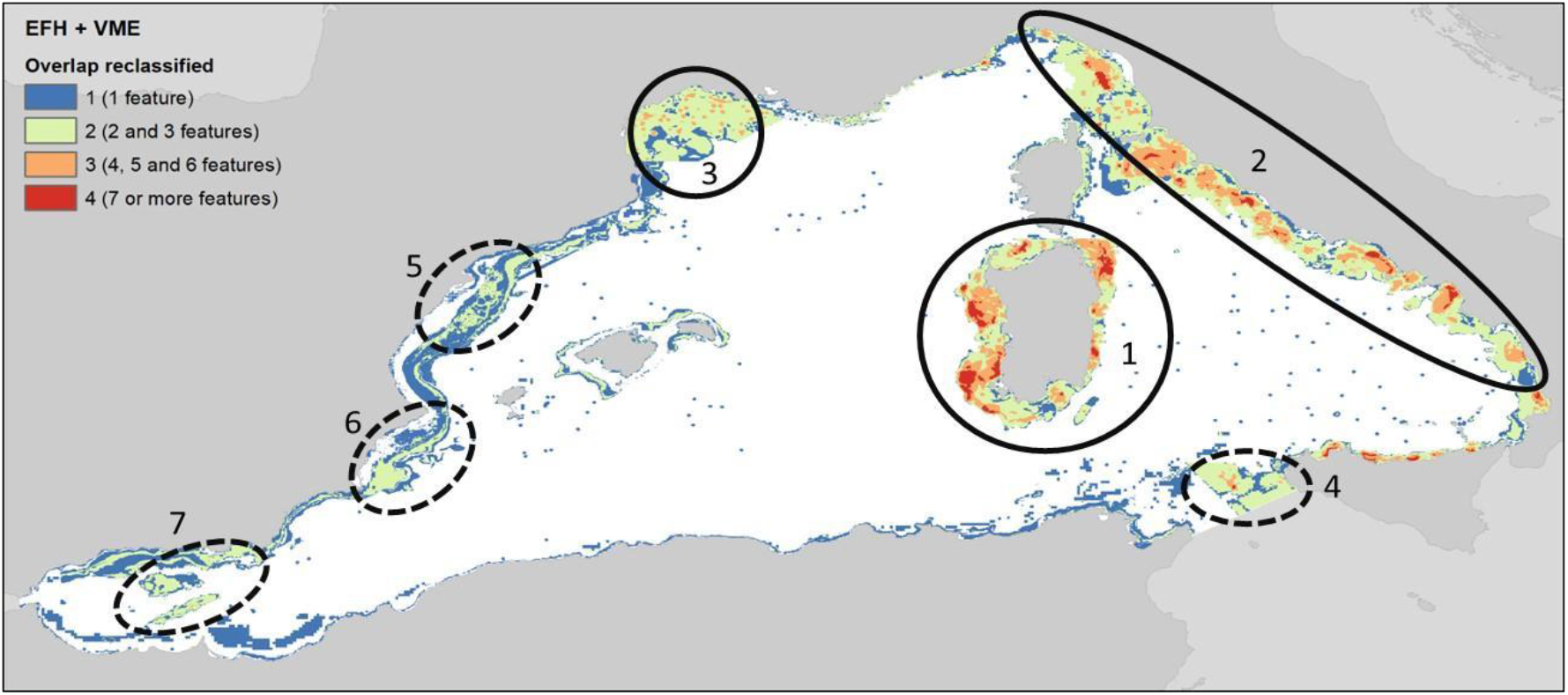
Multi-environmental value areas (MEVAs) map in the western Mediterranean Sea, with the seven regions of relatively high EFH + VME overlap highlighted.

### 3.2. Fishing pressure

High trawling effort occurs near the coast and over the continental shelves, with most intensity in the central areas of Spain and Italy. Relevant hotspots were found in the southern Ebro Delta (Spain) and Salerno and Northern Latium (Italy) (Figure 4 and Figure S5).

**Figure 4.**
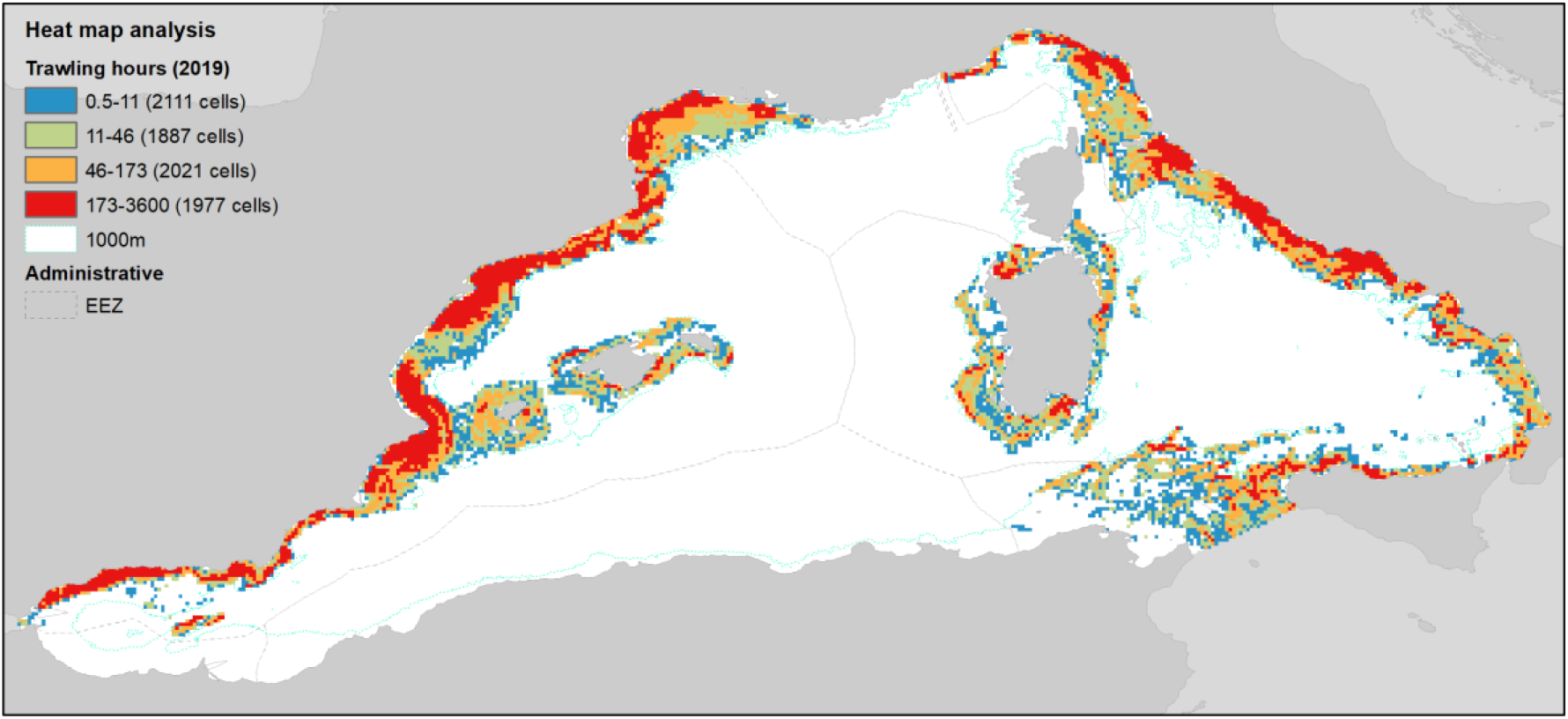
Fishing effort distribution classified by quartiles from low to high effort in the western Mediterranean Sea.

### 3.3. Priority Areas for Management (PAMs)

#### 3.3.1. Western Mediterranean Sea PAM

The PAM maps for the western Mediterranean show six candidate priority areas (Figure 5). The most relevant area surrounds Sardinia (Area 1), followed by the Italian Tyrrhennian waters (Area 2, with the most relevant subarea located in the north east), the Italian southern Tyrrhennian waters (Area 3, most notably in the northern part of the Strait of Sicily), the Gulf of Lions (Area 4), the Ebro Delta (Area 5) and around the Balearic Islands (Area 6). Other smaller areas are located in the Gulf of Alicante (Area 7) and in the Alboran Sea (Area 8).

**Figure 5.**
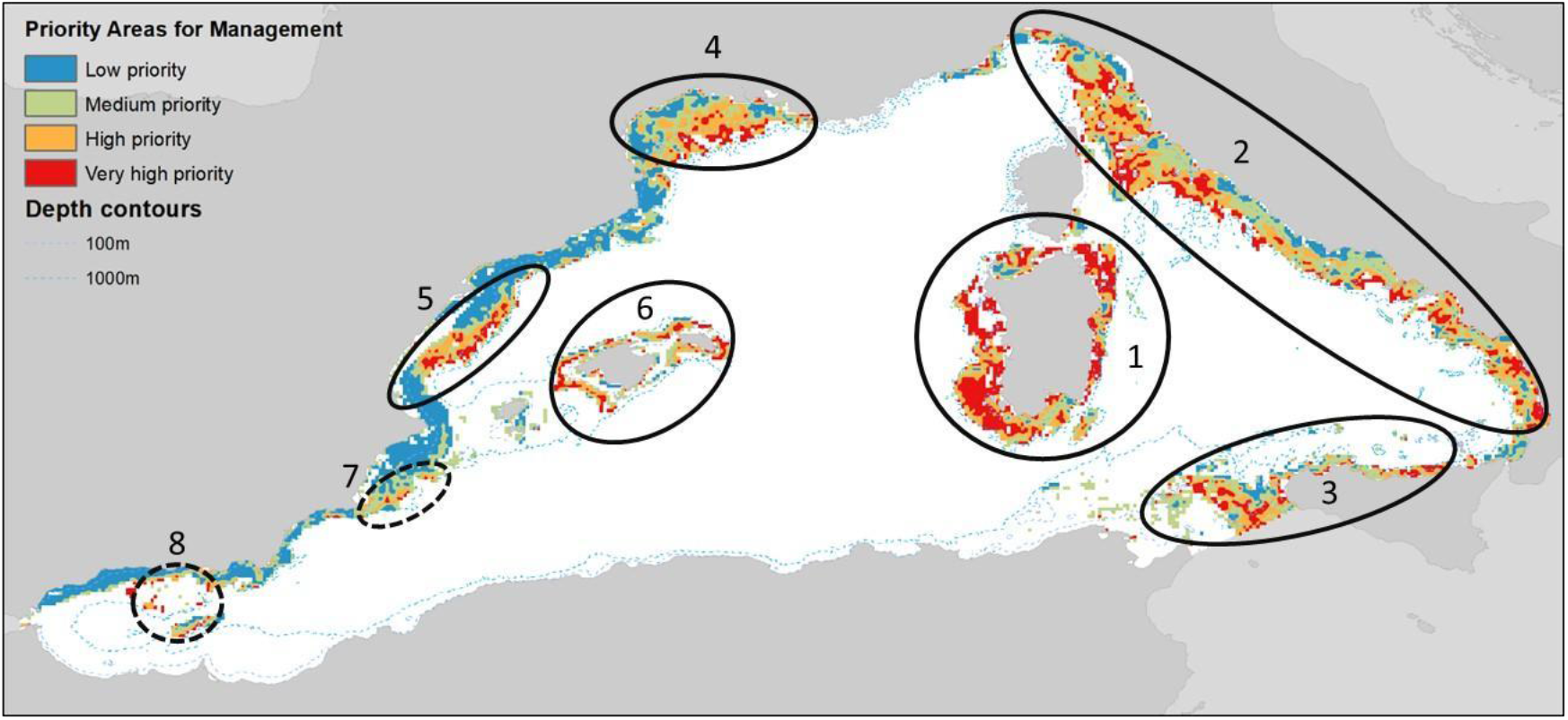
Priority Areas for Management in the western-Mediterranean Sea.

#### 3.3.2. EEZ-based PAMs

Mapping Priority Areas for Management for the EEZs of Spain, France and Italy, using only fishing pressure in each EEZ, showed slightly different patterns.

In the Spanish EEZ, the highest priority area for management can be discerned in the Ebro Delta, between Tarragona and Valencia, followed by a second potential area around the islands of Mallorca and Menorca. Some scattered smaller areas appeared in the Gulf of Alicante and the Alboran Sea (Figure 6a). In the northern part of the Spanish EEZ a potential area located next to French waters emerges, but when we analyzed the French EEZ (Figure 6b) it is clear that the highest priority from a French perspective lies in the central and Eastern part of the Gulf of Lions. In the Italian EEZ (Figure 6c), the area around Sardinia emerges as the highest PAM area, followed by areas in front of La Spezia and Civitavecchia.

**Figure 6.**
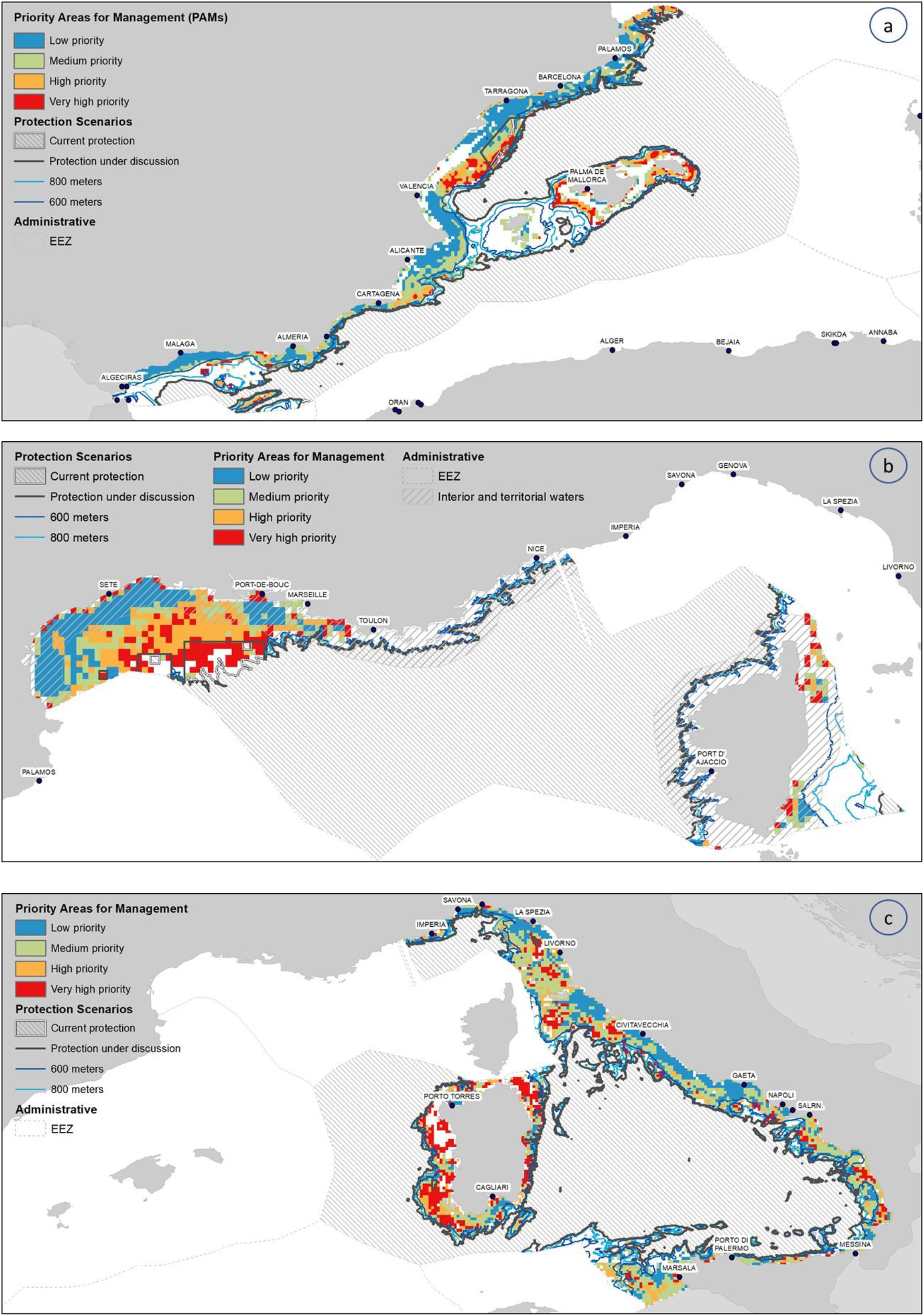
Priority Areas for Management for the EEZ of (a) Spain, (b) France and (c) Italy based on the distribution of trawling fishing pressure at each EEZ.

### 3.4. PAM protection under the four management scenarios

Spatial overlap between PAMs and protection scenarios across the entire western Mediterranean showed that current protection of PAMs is very low (Figure 7 and Table S6). However, if the discussed FRAs would be implemented (Figure 1 and Table 3), PAM protection would increase to 3% and 2.6% in high and very high priority areas, respectively. If we add to these protection figures the extension of the deep-sea FRA to 800 m of depth this protection would increase to 5% of high and very high priority areas. A further extension to 600 m depth would provide protection of approximately 10% of high and very high priority areas.

**Figure 7.**
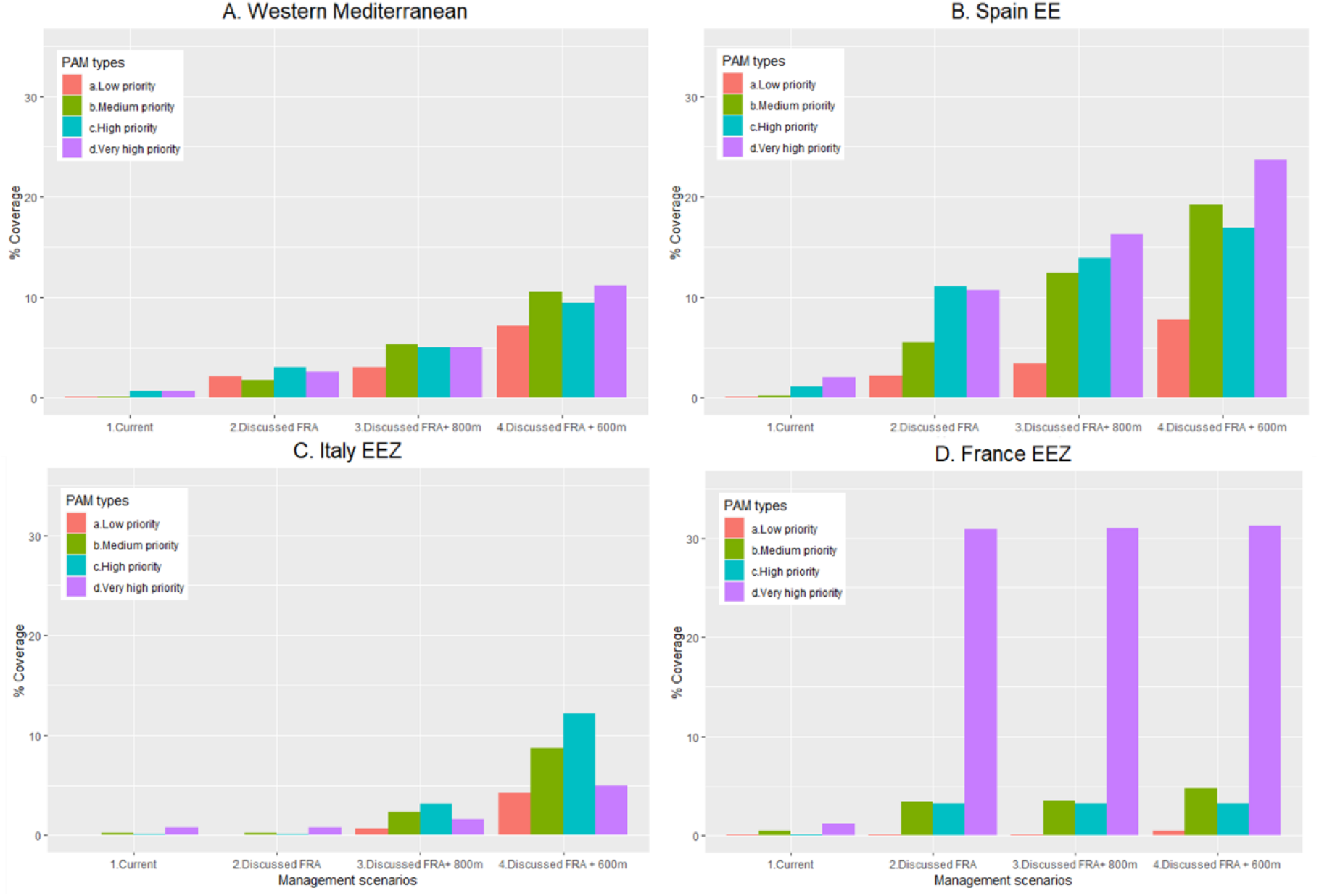
Percentage of PAMs overlapping with conservation figures in the a) Western Mediterranean, b) Spanish EEZ, c) French EEZ and d) Italian EEZ (Tables S6 and S7), excluding areas within 12 nautical miles.

When considering the PAM protection coverage per EEZ, the present situation is slightly better than the western average for Spain, while lower levels of protections than in the Spanish case are found for high and very high priority areas in France and Italy (Figure 7 and Table S6 and S7). Implementing the three discussed FRAs would notably increase the protection of PAMs in the French EEZ due to the closure of the FRA in the Gulf of Lions and the implementation of the Gulf of Lions Martí and Sète Canyons FRA, and in the Spanish EEZ due to the implementation of the Ebro Delta Margin FRA. Adding the deep-sea FRA expansions would be most relevant for Spain (for both high and very high priority areas), followed by Italy (most relevant for high priority areas), but would be less relevant for France.

## 4. Discussion

The creation of protected large marine areas for the recovery of Essential Fish Habitats and Vulnerable Marine Ecosystems conservation competes with other uses and planning interests, so there is an incentive to identify areas that simultaneously protect as many demersal commercial species and benthic communities as possible. We identified these areas, that we call “Multiple Ecological Values Areas” (MEVAs), throughout the western Mediterranean Sea.

Our results identified six areas with high EFH and VME overlap (Figure 3). Most of these areas are covered by the European Union “Multiannual plan for demersal stocks in the western Mediterranean Sea” and fall under European EEZ jurisdiction. The Strait of Sicily and the small MEVA in the Alboran Sea lie in both European and Non-European waters. The fact that we did not identify MEVAs in central North African waters may be due to ecological reasons but also to data paucity for this part of the Mediterranean (Coll et al., 2010).

### 4.1. From MEVAs to PAMs

Combining fishing intensity with MEVAs allowed for the identification of Priority Areas for Management: areas where adaptation measures are most feasible and ecologically relevant (Coll et al., 2015). Six main areas were identified as priority areas at the regional level (Figure 6): Sardinia, the northern eastern Italian coast, the northern Sicilian channel, the Gulf of Lions, the Ebro Delta and the Balearic Islands.

The PAM concept may be useful for refining and strengthening management proposals currently under discussion. For example, the GFCM “Recommendation GFCM/44/2021/5 on the establishment of a fisheries restricted area in the Gulf of Lion (geographical subarea 7) to protect spawning aggregations and deep-sea sensitive habitats, repealing eastern Gulf of Lion Recommendation GFCM/33/2009/1”, approved in 2021, established that, in 2023, the GFCM Scientific Advisory Committee on Fisheries will evaluate the implementation of the recommendation and advise the GFCM on further or alternative management measures addressing the overexploitation of demersal stocks, including a modification of the current closures, in time, geographical extension or gear use. The PAM concept provides additional insights to this evaluation.

Also, a complementary proposal for a FRA in the central area of the Gulf of Lion, the “Marti and Sète canyons FRA”, was submitted to the 2022 GFCM Subregional Committee for the Western Mediterranean in 2022 (FAO, 2022a). Both our regional and sub-regional findings support the Gulf of Lions as a priority area for protection.

In the Ebro Delta margin, a FRA proposal was endorsed by the SAC in 2021 as technically robust (FAO, 2021b). Our analysis, in coherence with other EFHs analysis in the area (DeLaHoz et al., 2018; Paradinas et al., 2022; Scientific Technical and Economic Committee for Fisheries, 2022a) and local knowledge (Bastari et al., 2022), supports this area as a priority area for management.

### 4.2. Evaluation of management measures

Our findings underpin that non-trawling areas currently provide a very limited coverage of PAMs at both the regional and sub-regional scale. In the western Mediterranean Sea, only 0.7% of the very high PAMs are off-limits to bottom trawling. Current levels of protection, with less than 2.5% of PAMs in all EEZs, indicate that the implemented areas are either too small, or are poorly placed, to provide the needed environmental protection (Scientific Technical and Economic Committee for Fisheries, 2022b). On the other hand, our analysis shows significant room to increase feasible protection of areas with ecologic relevance, even among protection measures that are already being considered at the GFCM.

The expansion of the deep sea FRA from 1000 meters to 800 or even 600 meters could be an effective measure to increase protection of high priority PAMs, both at regional (Figure 7 and Table S6) and sub-regional scales (Figure 7 and Table S7). At the western Mediterranean scale, the expansion scenarios analyzed could increase PAM protection sevenfold if the FRA was expanded from 800 meters depth, and sixteen-fold with an expansion to 600 meters.

The potential impact of the establishment of the proposed FRAs is also relevant at national levels. For example, in the French EEZ, the permanent closure of the Gulf of Lion FRA and the implementation of the proposed “Gulf of Lions Martí and Sete Canyons FRA” would increase protection of high priority areas by a factor 25. In the Spanish EEZ, the implementation of the “Ebro Delta Margin FRA” would increase PAM protection fivefold. In both cases, the full protection of these areas can be a relevant contribution to their respective conservation national commitments on the European Biodiversity Strategy for 2030 (European Commission, 2020), the Kunming-Montreal Global Biodiversity Framework framework (CBD, 2022) and the future European Nature Restoration regulation (European Commission, 2022). Last, our analysis strongly indicates that of all protection measures under discussion for Italian waters, the deep see FRA expansion is the only measure that will increase ecological protection.

It is important to highlight that we have not been able to analyze the impact of the “Alboran Sea Cabliers coral mound province FRA” (FAO, 2022a) due to lack of reliable fishing pressure data from North African countries. When new data becomes available our analyses can be redone to assess this proposal.

### 4.3. Scaling up the implementation of nature-based solutions

The potential of NbS for supporting fisheries sustainability and climate change adaptation is increasingly recognized (Key et al., 2021; Cooley et al., 2022; Seddon, 2022; Turner et al., 2022), yet further efforts are needed to better guide its operationalization (Riisager-Simonsen et al., 2022; Seddon, 2022).

Up to now, most marine NbS projects are small. In the western Mediterranean Sea, our MEVA and PAM analysis can provide relevant insights for the definition of scaled-up NbS towards sustainable harvesting practices, which are consistent with NbS principles (Cooley et al., 2022; Riisager-Simonsen et al., 2022). Our approach combines relevant criteria for NbS such as biodiversity net-gain and economic feasibility (Seddon, 2022; IUCN, 2020). Moreover, Mediterranean examples such as the Jabuka/Pomo Pit FRA (FAO, 2022b) or other smaller such as Roses non-take area in the North-Western Catalan Sea (Tuset et al., 2021) show that, if properly implemented, permanent FRAs and non-take areas in the Mediterranean can support the biomass recovery of priority species, thus contributing to fishing sustainability (Reimer et al., 2021).

Nevertheless, the provided information is just an entry point to the process of implementation of NbS. NbS must be designed, implemented, and monitored in close partnership with local communities rather than through top-down governance structures. Stakeholder participation in the definition process is key to gain the legitimacy to label the potential identified protected zones as or part of NbS (IUCN, 2020; Seddon, 2022).

### 4.4. Challenges and contributions of our approach

The identification of MEVAs and PAMs is complementary to previous analysis in the region, which focused either on the analysis of EFHs “hotspots” (Scientific Technical and Economic Committee for Fisheries, 2022a) or specific VMEs areas (Fanelli et al., 2021).

Nevertheless, our approach has also limitations that should be taken into consideration. First of all, EFH and VME data availability in the Mediterranean Sea is a challenge. EFH and VME data is incomprehensive and, even if some improvements have been achieved in the last years, out of date throughout the western Mediterranean, and is mostly lacking for the southern waters (Coll et al., 2010; FAO, 2022c; Ramírez et al., 2022). Our study represents a conservative view of MEVAs in the region and as new data becomes available the analyses presented can be updated.

Secondly, our results are also constrained by fishing pressure data availability. Global Fishing Watch uses Automatic Identification Systems (AIS) data when available, and Vessels Monitoring Systems (VMS) data for a limited number of countries. Currently the AIS system is mandatory for European fishing vessels exceeding 15 meters, while its use by the western Mediterranean non-EU fleets is very scarce. Recent GFW analysis for the whole Mediterranean estimated a substantial amount of fishing vessels that were not broadcasting AIS, with a significant fraction in Northern African waters^5^. While the implementation of a satellite-based VMS for all commercial fishing vessels exceeding 15 meters length system is mandatory for all vessels active in the GFCM area since 2012 (GFCM, 2009), key fishing nations in the area such as Algeria do not have yet a VMS system in place (GFCM, 2021b). According to the GFCM Authorized Vessel List (GFCM, 2022) the current Algerian fleet is composed of 506 active bottom trawlers in the western Mediterranean Sea, which is very similar to the Spanish or Italian bottom trawling fleet (565 and 541 vessels, respectively), and is much larger than the Moroccan and French trawling fleets (129 and 45 vessels, respectively). This indicates a significant data gap that limited our ability to analyze central North African waters.

Additionally, trawling pressure in the European Union waters has been officially decreasing since 2019, partially due to the application of the Multiannual plan for demersal fisheries in the western Mediterranean (Scientific Technical and Economic Committee for Fisheries, 2022c), which calls for an update of our analysis once post-COVID pandemic fishing effort data becomes available. Future MEVA and PAM analysis should also consider scenarios of environmental and socio-economic change. Adding future climatic data in MEVA and PAM analyses is an essential future step to identify areas that are likely to be affected by climate dynamics, which could serve to adapt the designation of FRAs with climate projections and plausible future uses in mind (Kyprioti et al., 2021, Pennino et al., 2020).

Despite these limitations, our results provide relevant input to the implementation of the GFCM 2030 Strategy in the western Mediterranean, and can be especially valuable taking into consideration that this organization is promoting a decentralized approach, so a western Mediterranean perspective is urgently needed (FAO, 2021b).

It is also important to note that we have not considered in our calculations a broad spectrum of spatial marine conservation schemes in the northern and western part of the region (see for example UNEP/MAP-SPA/RAC & MedPAN, 2022, or Natura 2000 sites) that are mostly linked with cetaceans, marine mammals and sea bird protection. However, as most of these schemes lack practical protection for demersal species and benthic habitats, they could not be considered in the MEVA and PAM analysis. Follow-up studies could broaden their scope for spatial conservation planning, considering multiple environmental values, stressors beyond fishing, and different priority species. Extending the MEVA and PAM concept, incorporating broader integrated ecosystem assessments of other ecosystem components, human activities and impacts, may offer valuable future contributions to conservation discussions and the assessment of good environmental status. As a final consideration, our scientific-based approach could be extended with local ecological knowledge and direct stakeholder involvement to identifying priority areas for management, following NbS principles.

## 5. Conclusions

There is a general consensus that severe anthropogenic pressures in the western Mediterranean are changing its characteristics at all ecological levels, and that urgent and targeted action is needed to recover exploited resources and to improve the marine environmental health. This is especially urgent for key commercial demersal species and benthic ecosystems, where many stocks are overfished (FAO, 2022d) and the current status of fragility of vulnerable marine ecosystems is clear (Bastari et al., 2022). The identification and prioritization of permanent fisheries restricted areas that simultaneously address the recovery of EFHs and the conservation of VMEs, and that can be easily implemented due to relatively low fishing pressure, can be a valuable contribution to address part of the problem, in line with the NbS schemes that are attracting an increasing scientific and political attention.

The systematic identification of six PAM areas based on the best publicly available data in the western Mediterranean basin aims at contributing to the political processes that are already in place at the international level, providing additional insights to ecosystem-based fisheries management strategies. Our analysis provides new integrated and complementary information to available data, and supports the implementation of marine spatial planning in the basin with evidence-based information.

Our analysis also shows that at the regional scale, recently established trawl no-take areas offer limited protection to PAMs, but that the implementation of the most extensive protection measures that are currently under discussion may offer significantly better protection.

Our analysis can provide useful information to scale-up marine NbS. The scientific-based approach described here offers the opportunity to explore areas that support biodiversity with lower presence of highly impacting fishing fleets. In practice, FRAs could be potential candidates to operationalize NbS when aligned with its standards and criteria, without overlooking the associated challenges, uncertainties and evidence gaps. Advancing on how, in practice, and where NbS could be implemented is essential for mainstreaming the use of NbS for supporting biodiversity, building resilience and climate change adaptation in the western Mediterranean Sea.

## Acknowledgements

Authors acknowledge partial funding from the EU Grant Agreement 101059877 – (GES4SEAS project) and EU GA 101059407 (MarinePlan project). The authors acknowledge the Spanish government through the ‘Severo Ochoa Centre of Excellence’ accreditation (CEX2019-000928-S). María D. Castro-Cadenas acknowledges the FPU grant of the Spanish Ministry of Universities (reference FPU2020/04852). The authors acknowledge the support of MedReAct and access to the database on VMEs from GFCM.

## Footnotes

1 https://globalfishingwatch.org/

2 (*Arrêté Du 20 Décembre 2019 Portant Modification de l’arrêté Du 28 Février 2013 Portant Adoption d’un Plan de Gestion Pour La Pêche Professionnelle Au Chalut En Mer Méditerranée Par Les Navires Battant Pavillon Français*, 2019; *Recommendation GFCM/44/2021/5 on the Establishment of a Fisheries Restricted Area in the Gulf of Lion (Geographical Subarea 7) to Protect Spawning Aggregations and Deep-Sea Sensitive Habitats, Repealing Eastern Gulf of Lion Recommendation GFCM/33/2009/1*, 2021);

3 (*Orden APA/423/2020, de 18 de Mayo, Por La Que Se Establece Un Plan de Gestión Para La Conservación de Los Recursos Pesqueros Demersales En El Mar Mediterráneo*, 2020);

4 (Decreto Di Attuazione Dell’art.6, Comma 1 Del D.M. N°13128 Del 31.12.2019 - Individuazione Delle Zone Vietate Alla Pesca Professionale Esercitata Con Gli Attrezzi “Rete a Strascico a Divergenti”, “Sfogliara Rapido”, “Reti Gemelle a Divergenti”, “Reti Da Traino Pelagiche a Coppia”, “Reti Da Traino Pelagiche a Divergenti” e “draghe Tirate Da Natanti (Ex Traino per Molluschi) Nelle GSA 9, 10 e 11 Ai Sensi Dell’art.11 Comma 2 Del Reg. (UE) N°1022/2019, 2020).

5 https://globalfishingwatch.org/data/radar-illuminated-ocean/

