## Supplementary material for "Identifying and prioritizing demersal fisheries restricted areas based on combined ecological and fisheries criteria: the western Mediterranean": Suplementary materials

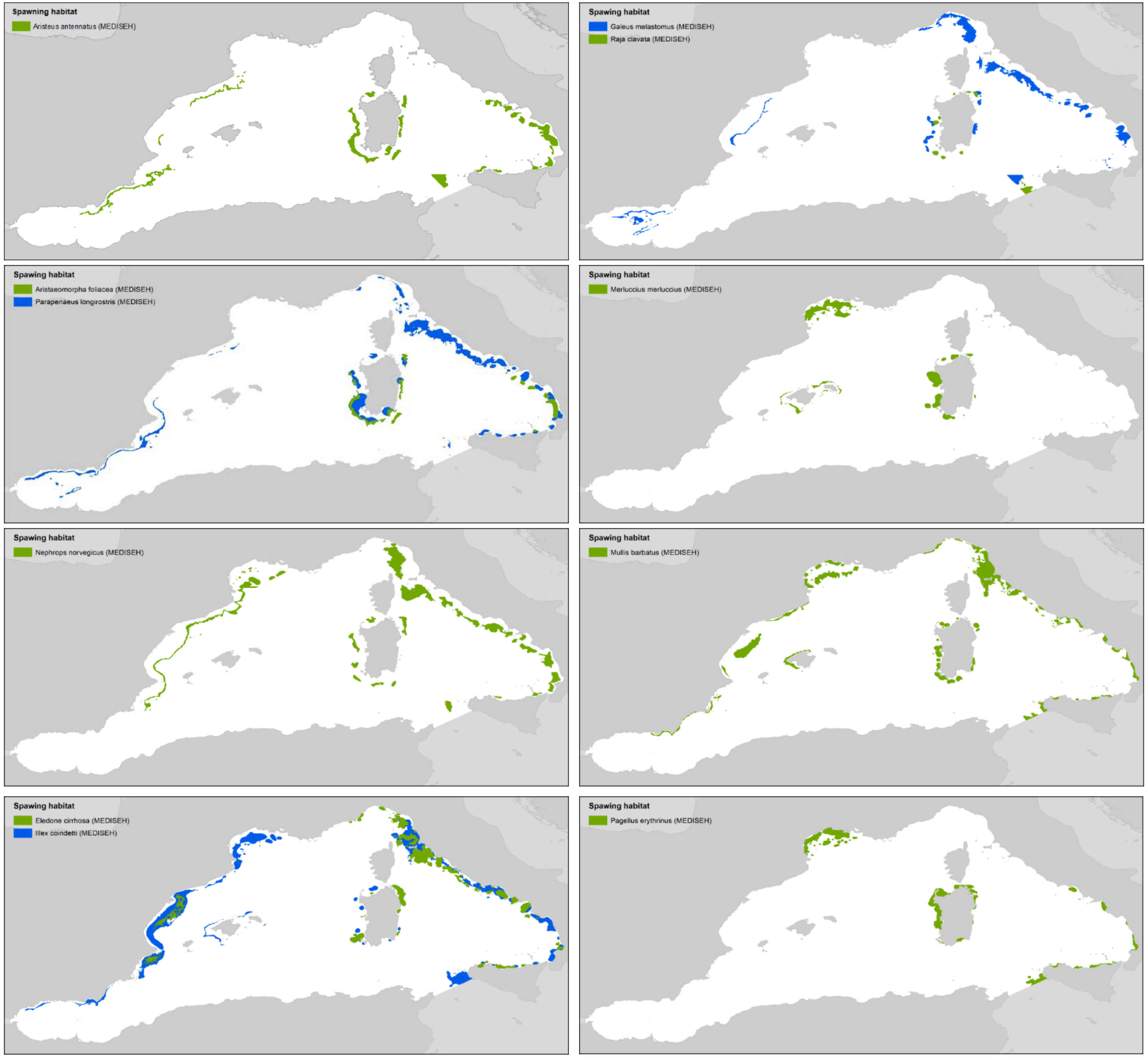


Figure S1. Available data for spawning areas of fish (*Merluccius merluccius, Mullus barbatus, Pagellus erythrinus*), elasmobranchs (*Raja clavata, Galeus melastomus*), cephalopods (*Eledone cirrhosa, Illex coindetii*), and crustaceans (*Aristaeomorpha foliacea, Parapenaeus longirostris, Nephrops norvegicus, Aristeus antennatus*).

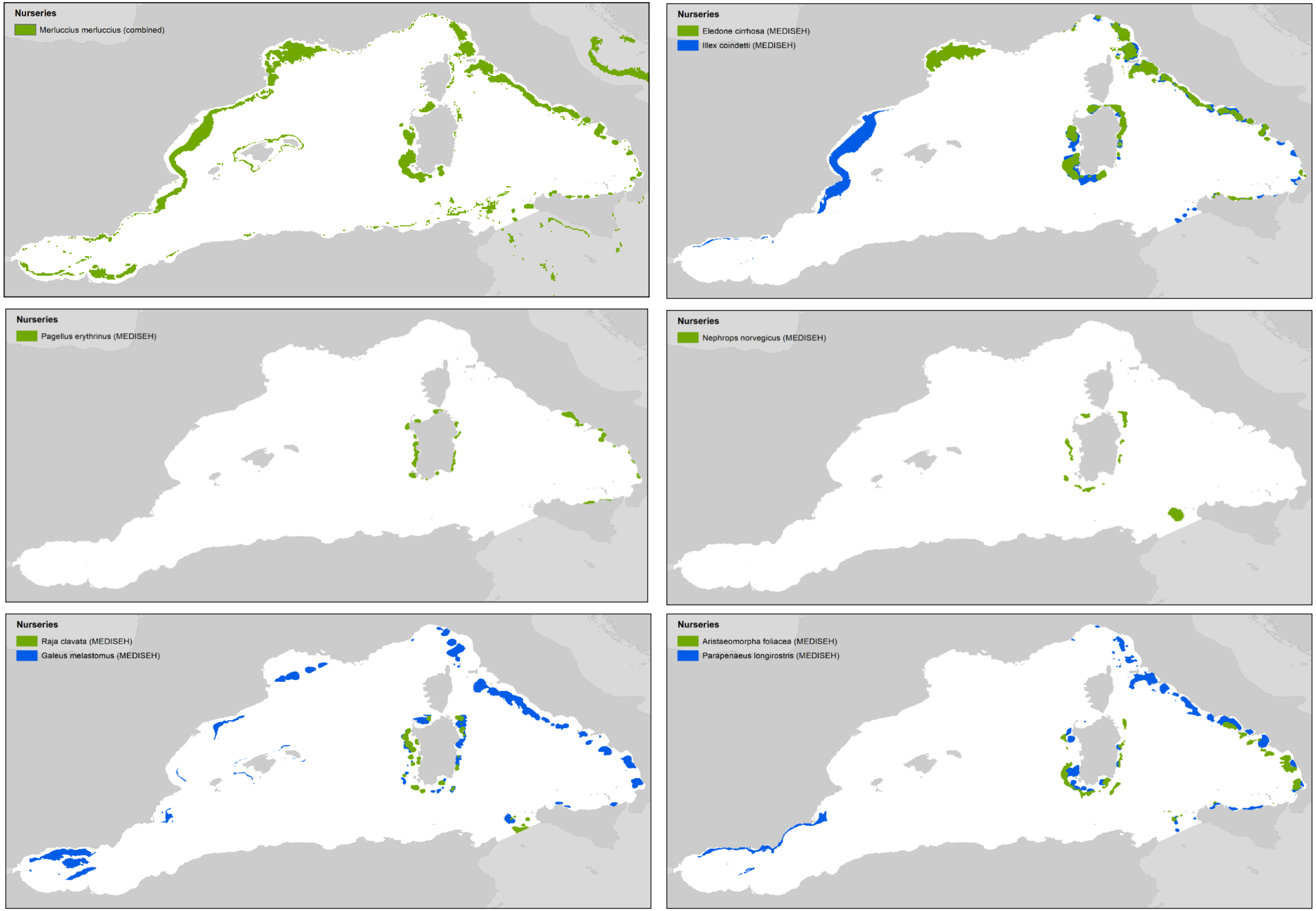


Figure S2. Available data for nursery areas of fish (*Merluccius merluccius, Pagellus erythrinus*), elasmobranchs (*Raja clavata, Galeus melastomus*), cephalopods (*Eledone cirrhosa, Illex coindetii*), and crustaceans (*Aristaeomorpha foliacea, Parapenaeus longirostris, Nephrops norvegicus*).

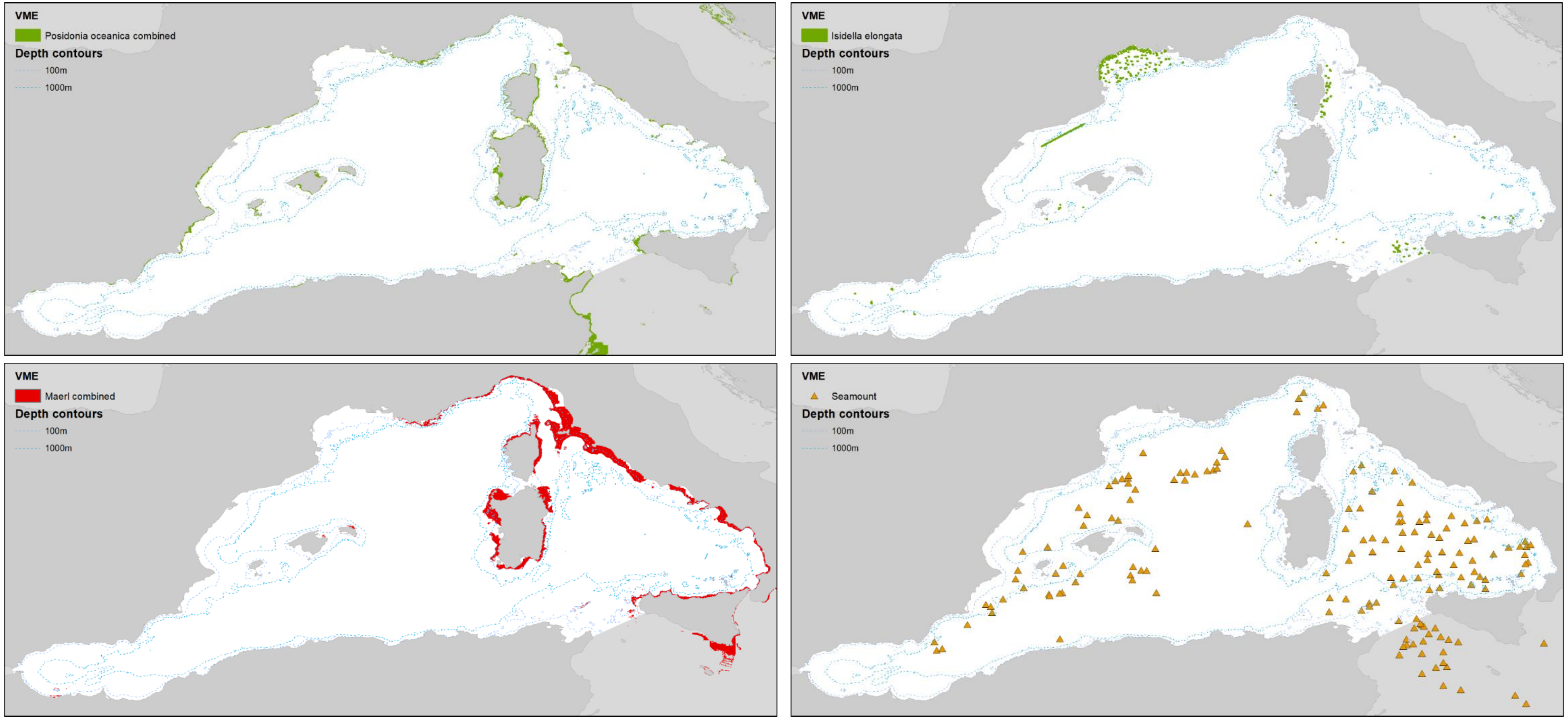


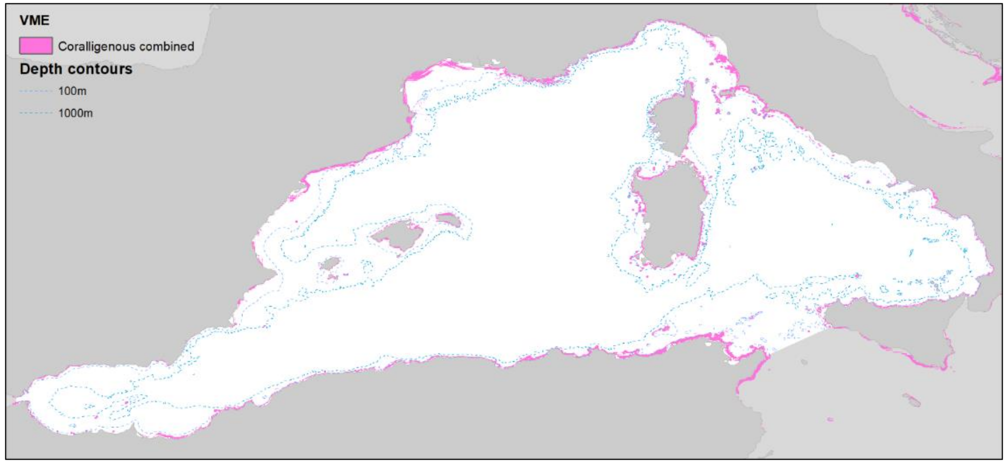


Figure S3. Available data for Vulnerable Marine Ecosystems (seagrass meadows, maërl beds, coralligenous communities, bamboo coral and seamounts).

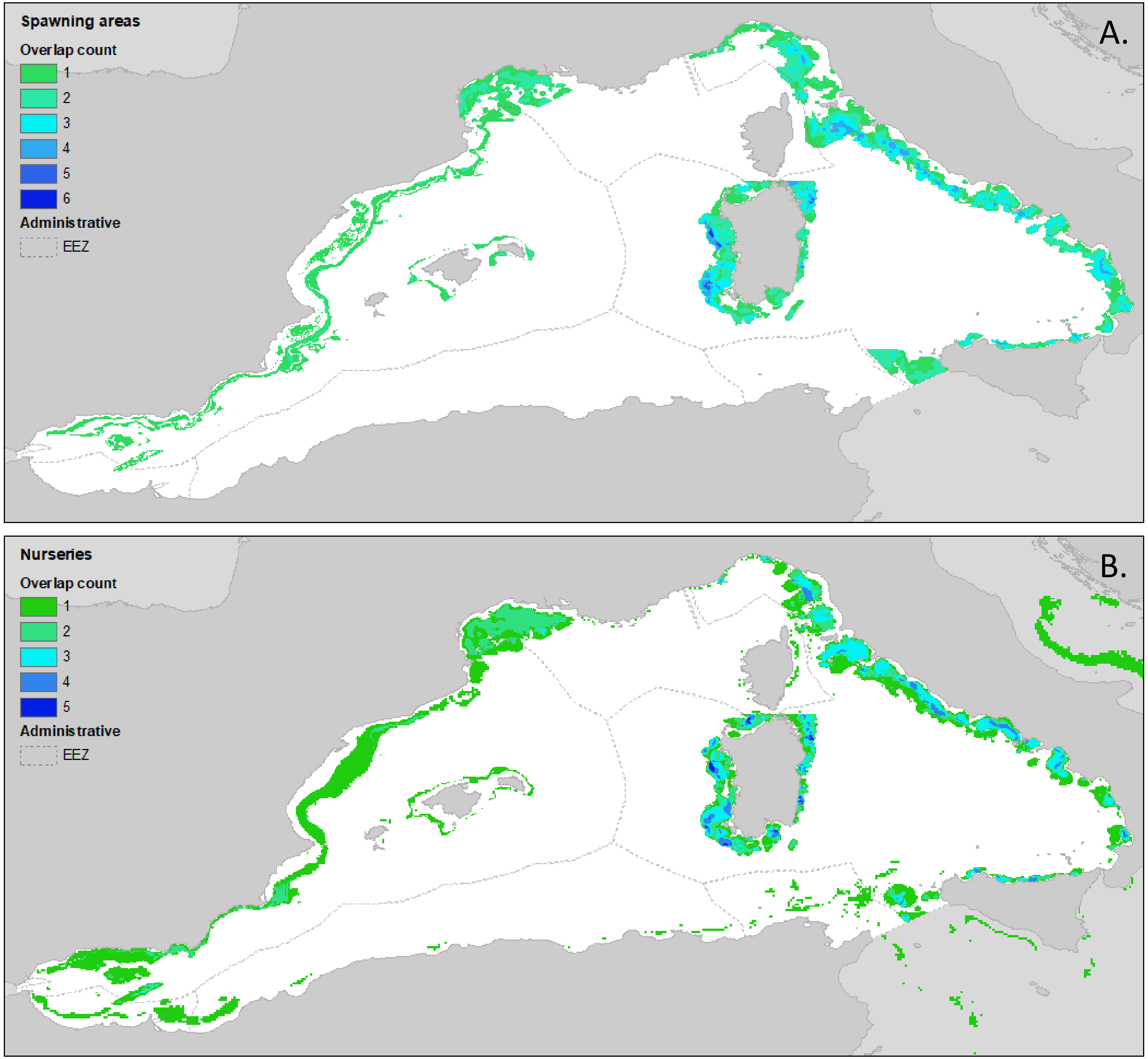


Figure S4. Available data for Essential Fish Habitats. a) Spawning areas data for the species *Aristeus antennatus*, *Eledone cirrhosa*, *Galeus melastomus*, *Merluccius merluccius*, *Mullus surmuletus*, *Nephrops norvegicus*, *Pagellus erythrinus*, *Parapenaeus longirostris; b)* Nursery areas data for the species *Eledone cirrhosa*, *Galeus melastomus*, *Merluccius merluccius*, *Nephrops norvegicus*, *Pagellus erythrinus*, *Parapenaeus longirostris*, *Raja clavata*

Figure S5. Bottom trawling fishing effort intensity in 2019 (source Global Fish Watch (GFW, 2022) coming from the Automatic Identification System (AIS)).

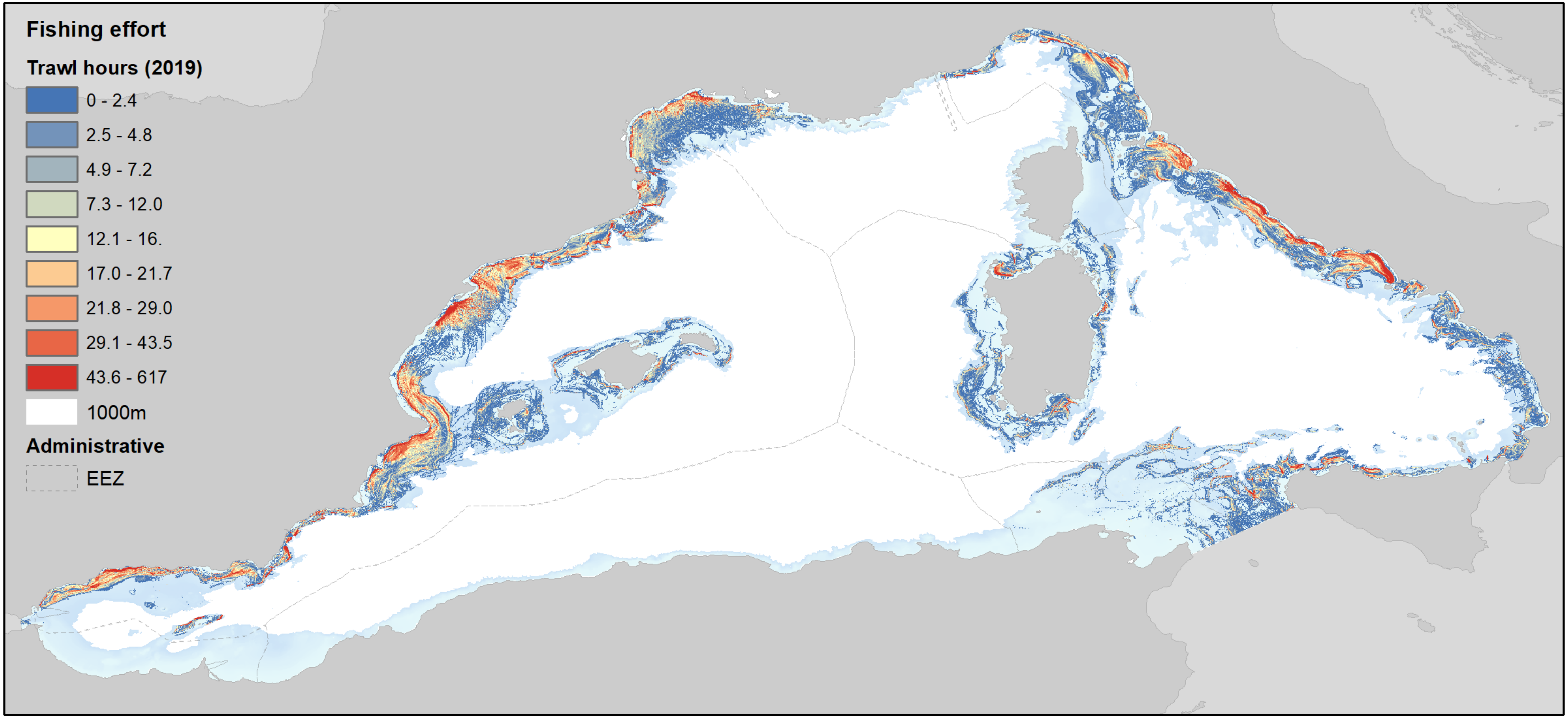

Table S6. Protection rates obtained from the overlapping analyses of conservation figures with Priority Areas Management at the western Mediterranean Sea scale, excluding areas within 12 nautical miles.

| Priority area coverage | Current implemented areas | Discussed FRA | Discussed FRA + 800m | Discussed FRA + 600m |
|---|---|---|---|---|
| Low priority | 0.08% | 2.11% | 3.05% | 7.11% |
| Medium priority | 0.13% | 1.74% | 5.31% | 10.57% |
| High priority | 0.70% | 3.03% | 5.03% | 9.47% |
| Very high priority | 0.70% | 2.58% | 5.04% | 11.15% |

Table S7. Protection rates obtained from the overlapping analyses of conservation figures with Priority Areas Management by EEZ, excluding areas within 12 nautical miles.

| | Spain EEZ | | | |
|---|---|---|---|---|
| Priority area coverage | Current implemented areas | Discussed FRA | Discussed FRA+ 800m | Discussed FRA+ 600m |
| Low priority | 0.09% | 2.24% | 3.41% | 7.78% |
| Medium priority | 0.24% | 5.51% | 12.43% | 19.20% |
| High priority | 1.11% | 11.03% | 13.93% | 16.95% |
| Very high priority | 2.07% | 10.70% | 16.32% | 23.67% |
| | France EEZ | | | |
| Priority area coverage | Current implemented areas | Discussed FRA | Discussed FRA + 800m | Discussed FRA+ 600m |
| Low priority | 0.08% | 0.08% | 0.09% | 0.49% |
| Medium priority | 0.52% | 3.43% | 3.48% | 4.76% |
| High priority | 0.09% | 3.20% | 3.20% | 3.22% |
| Very high priority | 1.26% | 30.84% | 30.99% | 31.26% |
| | Italy EEZ | | | |
| Priority area coverage | Current implemented areas | Discussed FRA | Discussed + 800m | Discussed + 600m |
| Low priority | 0.05% | 0.05% | 0.64% | 4.24% |
| Medium priority | 0.21% | 0.21% | 2.28% | 8.69% |
| High priority | 0.11% | 0.11% | 3.13% | 12.13% |
| Very high priority | 0.74% | 0.74% | 1.55% | 4.96% |
